# Recovery of proofreading-impaired SARS-CoV-2 reveals a mutator phenotype and an ExoN activity threshold for viability

**DOI:** 10.64898/2026.05.12.724615

**Authors:** Li He, Yuan-Wei Norman Su, Fushun Zhang, Ibrahim M. Moustafa, David W. Gohara, Chengjin Ye, Luis Martinez-Sobrido, Jamie J. Arnold, Craig E. Cameron, Yan Xiang

## Abstract

Coronaviruses (CoVs) replicate unusually large RNA genomes that necessitate proofreading by the 3′-to-5′ exoribonuclease (ExoN) formed by nonstructural proteins 14 (nsp14) and 10 (nsp10). Previous studies suggested that inactivation of the ExoN catalytic site in severe acute respiratory syndrome CoV 2 (SARS-CoV-2) is lethal, leaving unresolved whether the virus can tolerate impaired proofreading activity. Here, we investigated the functional requirement for ExoN in SARS-CoV-2 replication by combining a continuous fluorescence-based biochemical assay with an optimized single-bacmid reverse genetics system. Mutational analysis of residues involved in RNA binding or catalysis revealed graded effects on ExoN activity in vitro. Alanine substitution of Lys9, a residue positioned near the RNA-binding interface, did not reduce ExoN activity, whereas charge reversal at this position (K9E) impaired activity more strongly than alanine substitutions of the catalytic motif I residues D90 and E92 (D90A/E92A). Correspondingly, recombinant SARS-CoV-2 carrying K9A was readily recovered, whereas the D90A/E92A mutant was recovered only after an extended delay and K9E could not be rescued despite repeated attempts. The D90A/E92A mutant exhibited reduced replication while maintaining the engineered ExoN substitutions during serial passage. Deep sequencing of viral populations revealed a marked increase in genome-wide sequence variation in the D90A/E92A mutant, demonstrating a stable mutator phenotype. Together, these findings indicate that SARS-CoV-2 can tolerate substantial impairment of ExoN activity but depends on a minimal activity threshold for viability. This system provides a platform for defining how SARS-CoV-2 proofreading controls genome stability, viral fitness, and sensitivity to antiviral strategies that exploit reduced replication fidelity.

**Importance:** Coronaviruses have unusually large RNA genomes because they encode a proofreading enzyme that removes copying errors during replication. It has been unclear whether SARS-CoV-2 can survive when this proofreading function is strongly weakened, because earlier studies suggested that loss of the enzyme’s catalytic activity is lethal. We show that SARS-CoV-2 can tolerate substantial impairment of proofreading, but only when residual exonuclease activity remains above a minimal threshold. A virus with impaired proofreading replicates less efficiently and accumulates mutations across its genome, whereas a more severe defect prevents virus recovery. These findings clarify how coronavirus proofreading balances genome stability with viral fitness and provide a useful system for studying how reduced replication fidelity affects viral evolution, antiviral sensitivity, and attenuation. Defining this activity threshold may also help guide antiviral strategies that target coronavirus proofreading.

## Introduction

RNA viruses typically exhibit high mutation rates because their RNA-dependent RNA polymerases (RdRp) lack proofreading capability, which limits the genome size of most RNA virus families to below 15 kb [1]. Coronaviruses (CoVs) and some related virus families within the order *Nidovirales*, however, possess the largest known RNA genomes, which can exceed 30 kb [1]. This unusually large genome size has been attributed in part to the presence of a 3′-to-5′ exoribonuclease (ExoN) proofreading enzyme that enhances replication fidelity [1–3]. CoV ExoN removes misincorporated nucleotides or nucleotide analogs during RNA synthesis, thereby increasing replication fidelity and contributing to resistance against antiviral nucleotide analogs [4–6]. In addition to its role in proofreading, ExoN has also been implicated in evasion of host innate immune responses [7, 8].

CoVs are widely distributed in mammalian species and pose a persistent zoonotic threat to human health [9, 10]. Over the past two decades, three novel CoVs capable of infecting humans have emerged to cause epidemics: severe acute respiratory syndrome CoV (SARS-CoV), Middle East respiratory syndrome CoV (MERS-CoV), and severe acute respiratory syndrome CoV 2 (SARS-CoV-2), the causative agent of the COVID-19 pandemic [9, 11].

In SARS-CoV-2, ExoN is located in the N-terminal region of nonstructural protein 14 (nsp14), whereas the C-terminal region of nsp14 contains a guanine N7-methyltransferase domain involved in the 5′ capping of viral genomic and subgenomic mRNAs [12]. Nsp14 interacts with nsp10, which is essential for ExoN activity [13, 14]. The ExoN domain belongs to the DEEDh superfamily of exonucleases, which catalyze the excision of nucleoside monophosphates from RNA through a mechanism dependent on two divalent metal ions coordinated by five conserved residues organized in three catalytic motifs, motifs 1, II, and III [15–17]. Biochemical studies of CoV nsp14 have demonstrated that this enzyme degrades both single-stranded and double-stranded RNA (dsRNA), with a preference for the latter [16, 18, 19].

Interestingly, inactivation of ExoN through mutation of the canonical motif I residues produces different phenotypes across CoVs. Alanine substitution of the motif I DE residues in the betacoronaviruses mouse hepatitis virus (MHV) [20] and SARS-CoV [21] yields viable viruses that exhibit impaired replication fidelity, resulting in hypermutator phenotypes with approximately 15– to 20-fold increases in mutation rates. These mutants replicate in cell culture and in animal models but display increased sensitivity to mutagenic agents such as 5-fluorouracil [22, 23]. In contrast, equivalent mutations in the ExoN catalytic motif I have been reported to be lethal in other CoVs. Previous attempts to recover viable mutant viruses failed for the betacoronaviruses MERS-CoV and SARS-CoV-2 [24], the alphacoronaviruses HCoV-229E [2], transmissible gastroenteritis virus (TGEV) [7], and porcine epidemic diarrhea virus (PEDV) [25].

Despite the central role of ExoN in CoV replication, the degree to which SARS-CoV-2 can tolerate impaired ExoN activity remains unknown. Here, we combine quantitative biochemical analysis of defined nsp14 mutants with reverse genetics to examine how perturbations in RNA binding and catalysis affect ExoN function and viral replication. Our findings reveal a relationship between residual ExoN activity, viral viability, and genome stability. These results support a threshold model in which SARS-

CoV-2 can tolerate substantial impairment of proofreading activity but requires a minimal level of nsp14 ExoN function for productive replication.

## Results

### Development of a continuous fluorescence-based ExoN activity assay

Previous assays of ExoN activity have primarily relied on gel-based approaches that measure activity at limited time points. To enable continuous measurement of ExoN activity, we adapted a fluorescence-based assay originally developed for DNA exonucleases to monitor RNA degradation [26–28]. This assay is based on the fluorescent base analog 2-aminopurine (2AP), which exhibits increased fluorescence when released as a free nucleotide compared to 2AP-substituted single-stranded or double-stranded RNA. 2AP is an adenine analog that base pairs with uracil and has excitation and emission maxima at 310 and 380 nm, respectively (Fig. 1A). The assay relies on excision of 2AP from the 3′ end of a dsRNA primer-template substrate by ExoN, resulting in a time-dependent increase in fluorescence (Fig. 1B). The fluorescence intensity of free 2AP is linear over a broad concentration range (Fig. 1C) and is more than 10-fold higher than that of 2AP incorporated into RNA (Fig. 1D). When positioned at the terminal 3′ end of a dsRNA substrate opposite uracil, 2AP exhibited low fluorescence, providing a suitable substrate for monitoring ExoN activity (Fig. 1D). Addition of ExoN, with nsp10 present at a 10-fold molar excess, resulted in a concentration-dependent increase in fluorescence over time (Fig. 1E).

**Figure 1.**
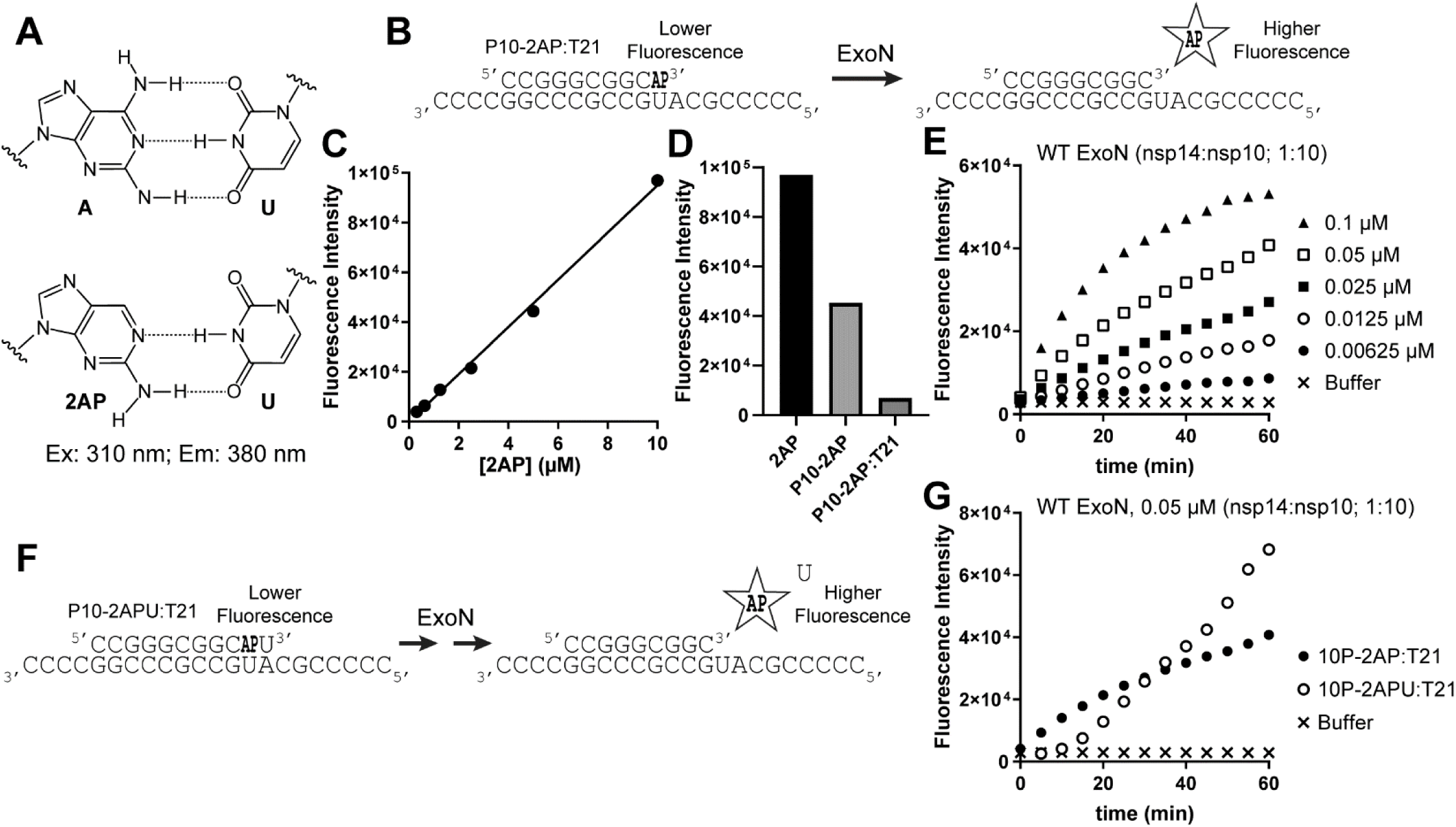
A 2-aminopurine (2AP) fluorescence-based assay to assess SARS-CoV-2 ExoN activity. **(A)** Chemical structure of the A:U and 2AP:U basepair. 2-aminopurine is a fluorescent analog with an excitation maximum of 310 nM and emission maximum of 380 nm. The fluorescence of 2AP is quenched when basepaired. **(B)** Schematic of ExoN assay. 2AP fluorescence is low when present in dsRNA, but when ExoN excises the 2AP nucleoside there is an increase in fluorescence. **(C)** Fluorescence intensity of the 2AP nucleoside from two-fold serial dilution (0.3 to 10 µM). **(D)** Comparison of 2AP fluorescence intensity between equimolar concentrations of 2AP nucleoside, 10P-2AP single-stranded RNA, and 10P-2AP:T21 double-stranded RNA. Note, the large difference in fluorescence intensity between the different forms of 2AP. **(E)** Kinetics of ExoN activity by monitoring the change in 2AP fluorescence intensity using varying concentrations of the ExoN nsp10:nsp14 complex. The ratio of nsp10 to nsp14 was kept constant at 10:1. **(F)** Schematic of exoribonuclease assay when 2AP is at the penultimate position of the RNA primer strand, 10P-2APU:T21. **(G)** Comparison of the kinetics of exoribonuclease activity with 2AP at either the ultimate or penultimate position using 10P2AP:T21 and 10P-2APU:T21 as substrates with a fixed concentration of ExoN WT.

We also evaluated a second 2AP-containing dsRNA in which 2AP was present at the penultimate position (Fig. 1F). For substrates such as this type, the rate-limiting step (*k*_cat_) in the steady state is most likely dissociation of ExoN from the substrate it just acted on to another substrate. If the rate of hydrolysis of two nucleotides is on par with the rate of dissociation, the reaction should exhibit a lag followed by a linear accumulation of product, as observed (Fig. 1G). Mutations that destabilize the ExoN-RNA complex would be expected to exhibit *k*_cat_ values greater than that of wild type (WT). If the rate of hydrolysis is reduced to a value below the rate of dissociation, then differences between the two substrates would be expected. For the 2AP-terminated substrate, a linear accumulation of fluorescence at a lower rate than WT would be expected. For the 2AP-recessed substrate, a longer lag followed by linear accumulation of product would be expected. If the ExoN-RNA becomes 10-fold or so more stable, then both substrates would exhibit *k*_cat_ values lower than wild type and loss or shortening of the lag using the 2AP-recessed substrate.

### Identification of nsp14 mutations that differentially impair ExoN activity

Structural studies of the SARS-CoV-2 nsp10-nsp14-RNA complex identified a basic RNA-binding surface adjacent to the ExoN active site [16, 18]. K9, K61, and K139 are located within or near this interface [18] (Fig. 2A), and alanine substitution of these residues reduced ExoN-mediated dsRNA degradation to varying degrees in a gel-based endpoint assay [18]. To compare the effects of perturbing RNA-binding or the catalytic-site on ExoN function, we expressed and purified nsp14 proteins carrying K9A, K9E, K61A, K139A, or the catalytic motif I substitutions D90A/E92A and assayed them using a dsRNA substrate containing 2AP at the penultimate position.

**Figure 2.**
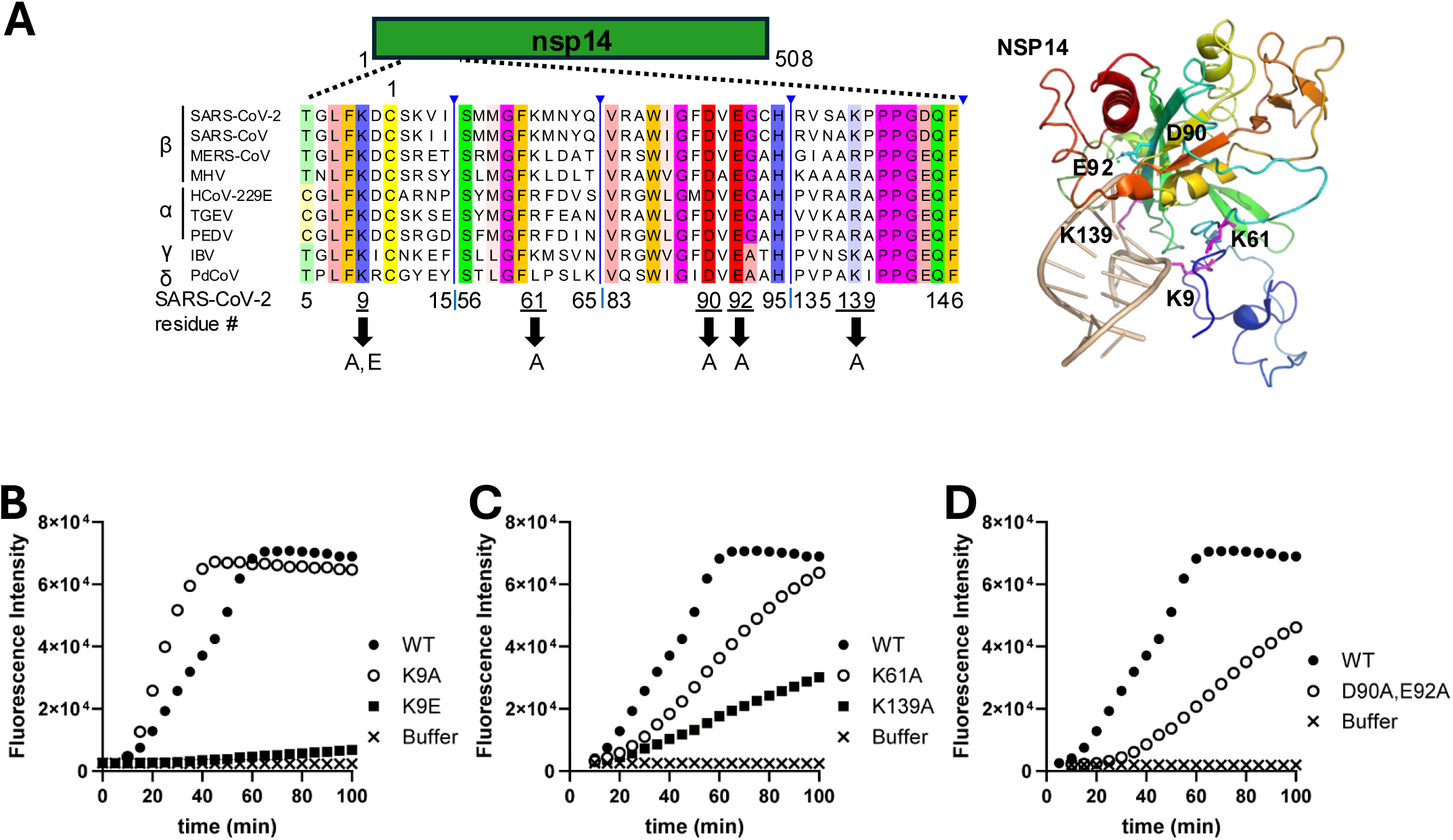
Nsp14 mutations differentially impair SARS-CoV-2 ExoN activity. **(A)** A partial multiple sequence alignment of ExoN from representative CoVs, highlighting residues targeted for mutagenesis in this study. These residues are also indicated and shown as sticks in the structure of SARS-CoV-2 nsp14 in complex with double-stranded RNA (PDB: 7N0B). **(B-D)** Comparison of ExoN activity between nsp14 WT and mutants using 10P-2APU:T21 dsRNA substrate. **(B)** WT, K9A, and K9E. **(C)** WT, K61A, and K139A. **(D)** WT and D90A/E92A.

The nsp14-K61A and –K139A proteins displayed reduced activity relative to WT to different extents (Fig. 2C), consistent with decreased stability of the ExoN-RNA complex. In contrast, the nsp14-K9A mutant exhibited a higher apparent rate of hydrolysis than WT, whereas nsp14-K9E was severely impaired (Fig. 2B). These results suggest that substitution of K9 with glutamate introduces electrostatic repulsion that destabilizes the ExoN-RNA complex and prevents the RNA substrate from adopting a conformation optimal for catalysis.

The nsp14-D90A/E92A protein exhibited the expected lag phase associated with a substantially reduced catalytic rate but showed substantially more activity than nsp14-K9E (Fig. 2C). The residual activity of nsp14-D90A/E92A likely reflects compensation by the extensive Mg²⁺ coordination network, which may partially mitigate the loss of the D90 and E92 catalytic residues, as suggested by molecular dynamics simulations (Fig. S1). These results indicate that these SARS-CoV-2 nsp14 mutations impair ExoN activity to different degrees, with K9A having little impact, D90A/E92A causing substantial catalytic impairment, and K9E producing the most severe defect.

### Alanine substitutions of lysine residues near the RNA-binding site are well tolerated in an attenuated SARS-CoV-2

The graded biochemical effects of the nsp14 mutations allowed us to test whether SARS-CoV-2 viability requires an ExoN activity threshold. We therefore attempted to rescue recombinant SARS-CoV-2 with various nsp14 mutations. We first introduced the nsp14-K9A, –K61A, and –K139A mutations individually into an attenuated SARS-CoV-2 lacking the open reading frames (ORFs) 3a and 7b [29] (Fig. 3A). This attenuated background was initially selected to potentially permit studies at a reduced biosafety level.

**Figure 3.**
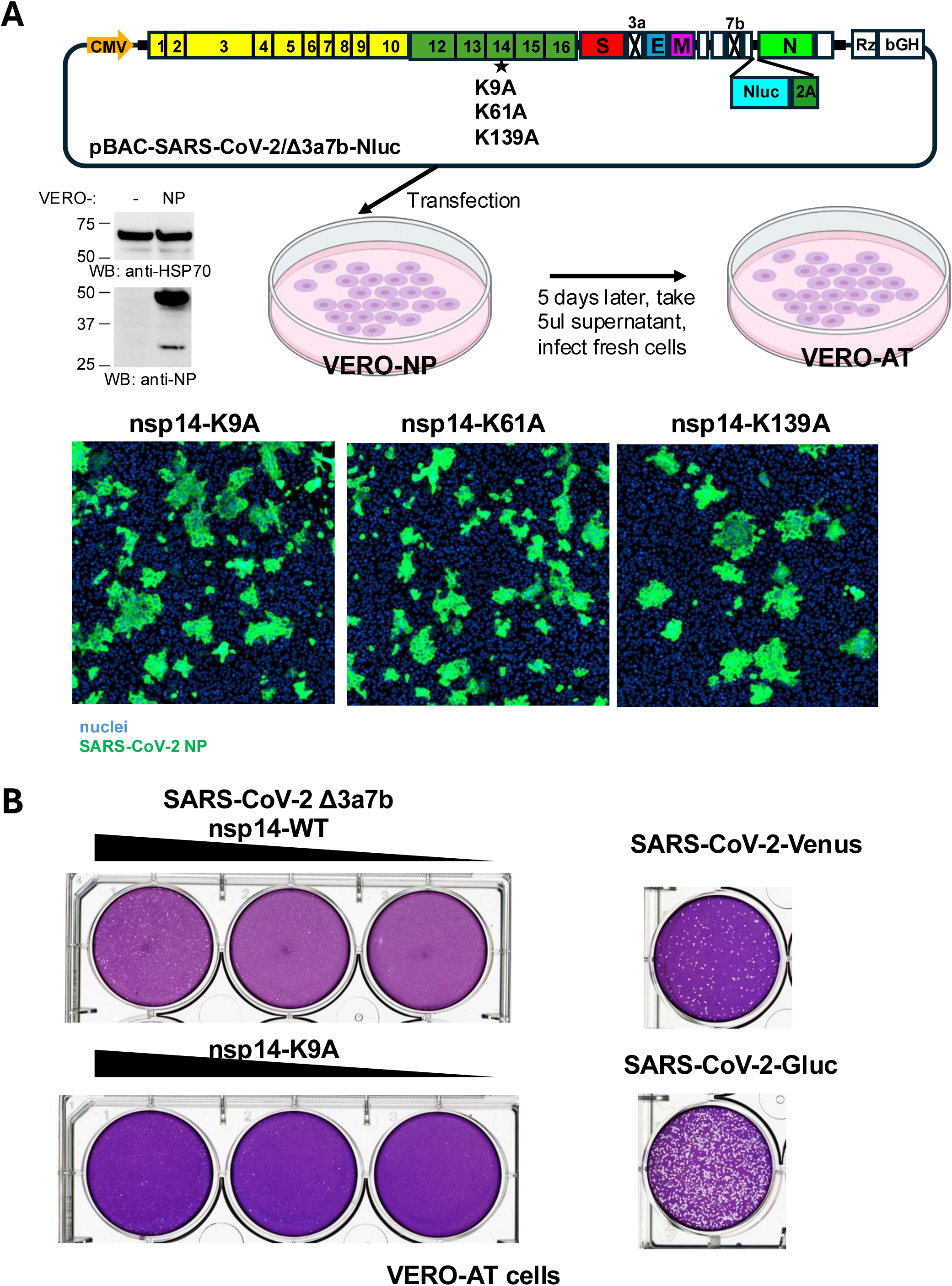
Alanine substitutions of lysine residues near the RNA-binding site are well tolerated in an attenuated SARS-CoV-2. **(A)** Generation of nsp14 mutant viruses. The SARS-CoV-2 genome in the bacmid contains deletions of ORF3a and ORF7b and is under the control of the CMV promoter. A Vero cell line expressing the SARS-CoV-2 N protein (Vero-NP) was engineered, and N expression was confirmed by immunoblotting. Bacmid DNA carrying engineered nsp14 mutations (K9A, K61A, or K139A) was transfected into Vero-NP cells, and supernatants were used to infect Vero-AT cells. Infected cells were analyzed 1 day later by immunofluorescence staining for N (green) and nuclei (blue). **(B)** Plaque morphology of recombinant viruses on Vero-AT cells. The triangle indicates three 10-fold serial dilutions of the viruses. Viruses in the Δ3a7b background (nsp14-WT or nsp14-K9A) produced smaller plaques than nonattenuated viruses expressing either Venus or Gluc.

To improve the efficiency of recombinant SARS-CoV-2 generation, we used a single-bacmid-based, self-launching reverse genetics system together with an engineered Vero cell line expressing the SARS-CoV-2 N protein (Vero-NP) (Fig. 3A), as N is known to facilitate virus recovery. To enable precise mutagenesis of the SARS-CoV-2 genome within the large bacmid, we employed *en passant* bacmid engineering technique based on lambda Red-mediated homologous recombination in bacteria. Following transfection of bacmid DNA into Vero-NP cells, supernatants were harvested and used to infect Vero cells engineered to express human ACE2 and TMPRSS2 (Vero-AT cells).

Immunofluorescence staining for N protein revealed clusters of infected cells of similar size (Fig 3A), indicating that all three mutants are viable.

To further assess viral growth, the nsp14-K9A mutant viruses was harvested, and its plaque morphology on VERO-AT cells was examined (Fig. 3B). The nsp14-K9A mutant formed very small plaques similar in size to those of the parental rSARS-CoV-2 Δ3a7b-Nluc virus. In contrast, recombinant SARS-CoV-2 containing the full complement of viral genes, such as rSARS-CoV-2/Venus and rSARS-CoV-2/Gluc, formed substantially larger plaques, consistent with a role for ORF3a and ORF7b in efficient virus replication.

### The nsp14-D90A/E92A mutation produces viable SARS-CoV-2 with reduced replication efficiency

We next evaluated nsp14 mutants that showed greater reductions in ExoN activity in vitro: nsp14-D90A/E92A and nsp14-K9E. We introduced these mutations into a bacmid containing the full-length SARS-CoV-2 cDNA with an additional mCherry-Nluc fusion reporter (rSARS-CoV-2/mCherry-Nluc) [30], allowing fluorescent protein expression to serve as a convenient readout for recombinant virus recovery. Following transfection into Vero-NP cells, rescue of the nsp14-WT and –K9A viruses was detected by mCherry expression within approximately 3 to 6 days. In contrast, the nsp14-D90A/E92A mutant required approximately two weeks before mCherry-positive cells became detectable (Fig. 4A). Despite repeated attempts, the nsp14-K9E mutant could not be rescued, even three weeks after transfection.

**Figure 4.**
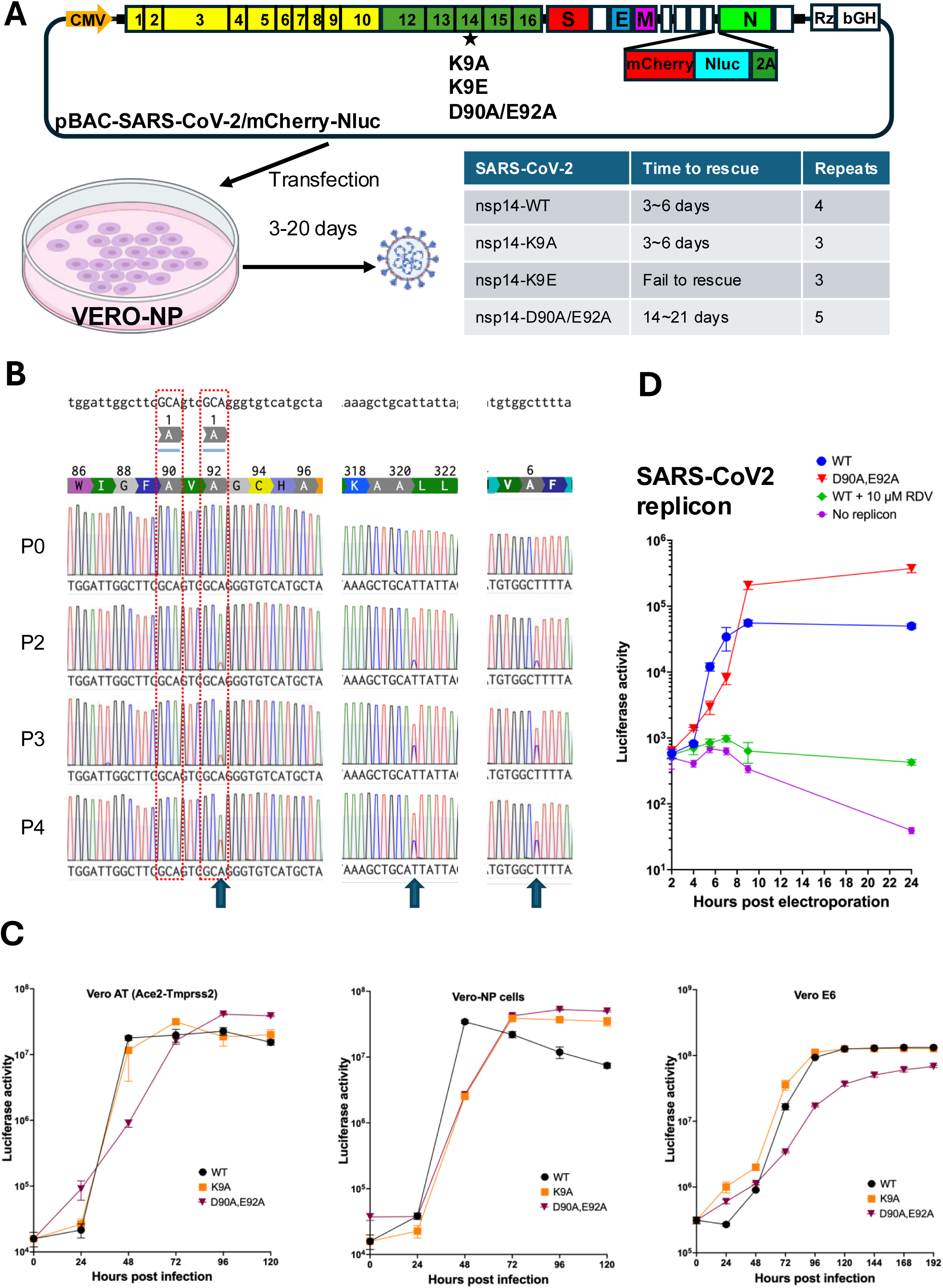
The nsp14-D90A/E92A mutation produces viable SARS-CoV-2 with reduced replication efficiency. **(A)** Generation of nsp14 mutant viruses. The bacmid contains the Washington strain SARS-CoV-2 genome carrying dual reporters (mCherry and Nluc) linked to N through a 2A sequence. Bacmid DNA carrying engineered nsp14 mutations was transfected into Vero-NP cells, and recovery of infectious virus was monitored by mCherry expression for up to 21 days. The table shows the time required for rescue and the number of independent rescue attempts for each virus. nsp14-K9A was readily rescued, nsp14-K9E was not recoverable, and nsp14-D90A/E92A exhibited delayed rescue. **(B)** Sequencing of nsp14 amplicons from different passages of the nsp14-D90A/E92A mutant virus. Arrows indicate positions showing mixed nucleotide peaks. **(C)** Replication kinetics of mutant viruses in different cell lines. Vero-AT, Vero-NP, and Vero E6 cells were infected with nsp14-WT, nsp14-K9A, or nsp14-D90A/E92A at an MOI of 0.1, and luciferase activity in culture supernatants was measured at the indicated time points after infection as a proxy for virus production. **(D)** SARS-CoV-2 replicon assay. SARS-CoV-2 replicon RNA was generated by in vitro transcription and electroporated into 293T cells. Luciferase activity was measured at the indicated time points. Remdesivir (10 μM, RDV) was included in one sample as a control.

We initially assessed the genetic stability of the mutant viruses by isolating viral RNA from different passages and sequencing a 1.6 kb amplicon spanning the nsp14 region. All passages examined, including P0, P2, P3, and P5, retained the engineered mutations (Fig. 4B). However, we also observed a progressive increase in synonymous substitutions at three positions within the amplicon, suggesting an elevated mutation rate.

To compare replication properties, we infected cells with nsp14-WT, –K9A, or – D90A/E92A viruses and measured luciferase activity in the cell culture supernatant as a proxy for viral release (Fig. 4C). The nsp14-K9A mutant displayed replication kinetics similar to those of the WT virus in all cell lines tested. In contrast, the nsp14-D90A/E92A mutant exhibited reduced replication in Vero E6 cells, although no significant reduction in luciferase activity was observed in Vero-AT or Vero-NP cells.

We next assessed viral growth by plaque assays in Vero-AT cells and in BHK cells engineered to express human ACE2 (BHK-hACE2), as the latter cell line allowed SARS-CoV-2 to form larger plaques. The nsp14-WT and –K9A viruses formed plaques of similar size in Vero-AT cells, whereas the nsp14-D90A/E92A mutant produced much smaller plaques that were barely visible (Fig. 5). The nsp14-D90A/E92A mutant formed larger plaques in BHK-hACE2 cells, but their size remained substantially smaller than those by the nsp14-WT virus.

**Figure 5.**
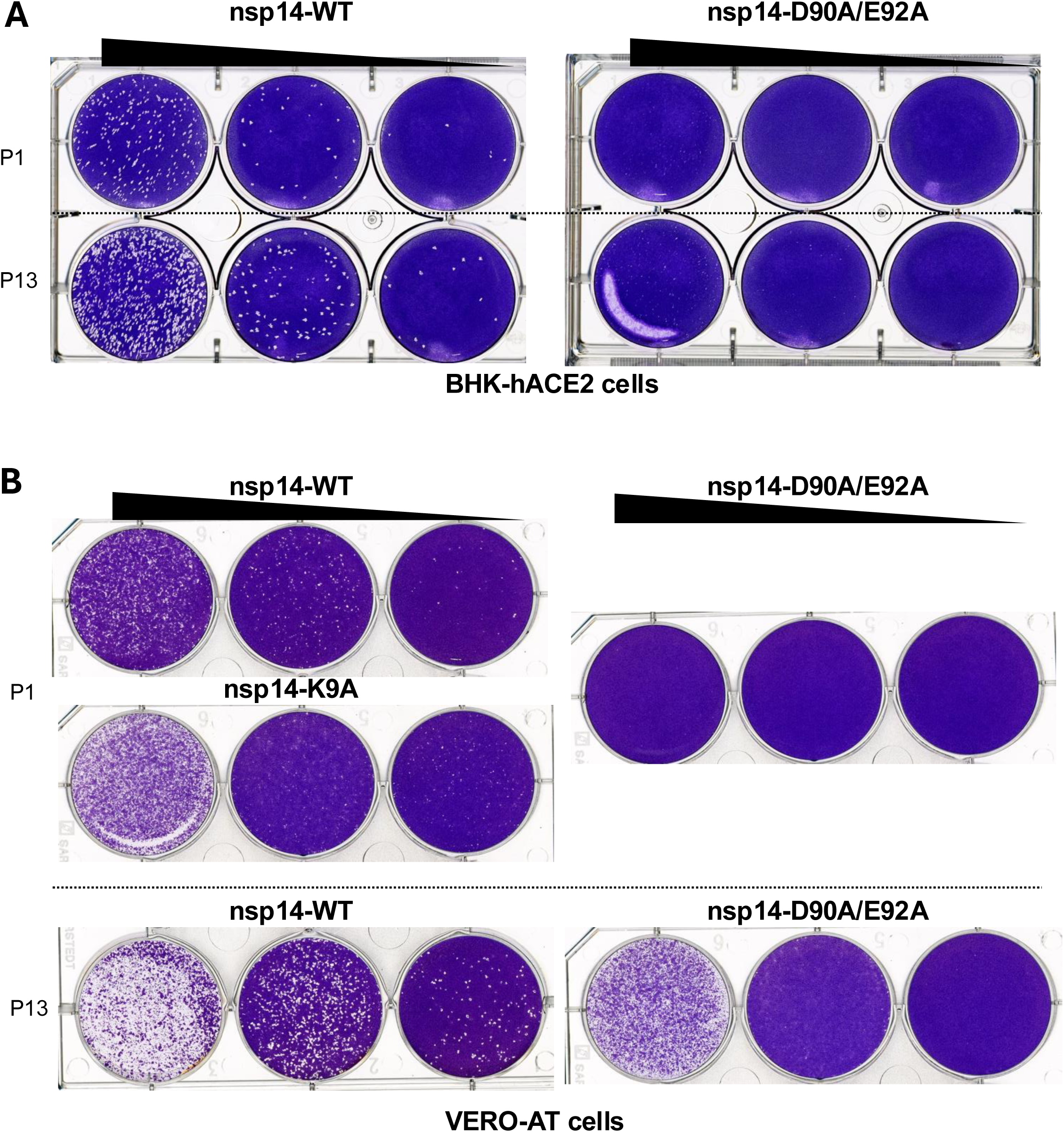
Plaque morphology of recombinant SARS-CoV-2 nsp14 mutants. Representative plaque assays of the indicated recombinant SARS-CoV-2 nsp14 mutants performed on BHK cells engineered to express human ACE2 and on Vero-AT cells. The triangle indicates three 10-fold serial dilutions of the viruses.

To determine whether the nsp14-D90A/E92A mutation affects viral RNA synthesis, we also introduced the mutation into a SARS-CoV-2 replicon (Fig. 4D). In vitro-transcribed replicon RNA was electroporated into HEK293T cells, and RNA production was measured indirectly by replicon-encoded luciferase activity. The nsp14-D90A/E92A replicon exhibited luciferase activity comparable to that of the nsp14-WT replicon, indicating that this mutation does not cause a major defect in replicon reporter expression under these conditions.

Together with the biochemical data, these rescue phenotypes support a threshold model for SARS-CoV-2 ExoN function. K9A, which retains robust activity in vitro, was readily recovered. D90A/E92A, which substantially impairs catalytic activity but does not abolish detectable activity in our assay, was recoverable only after a prolonged delay and showed reduced replication. In contrast, K9E, which produced the most severe biochemical defect, could not be rescued.

### The nsp14-D90A/E92A mutation reduces SARS-CoV-2 replication fidelity

To determine the impact of ExoN impairment on viral replication fidelity, we serially passaged the nsp14-WT and nsp14-D90A/E92A viruses in Vero-NP (Fig. 6) or Vero-AT cells (Fig. S2) and performed deep sequencing analysis of viral genomic RNA. Read depth analysis showed that all samples maintained comparable sequencing coverage across the viral genome at different passages (Supplemental data 1). One exception was the mCherry-Nluc reporter region, which was progressively lost in nsp14-WT virus during later passages (Fig. 6B and S2B). This loss was not observed for the nsp14-D90A/E92A mutant during passage in Vero-NP cells. However, deletions of ORF7b and ORF8 were detected in later passages of the nsp14-D90A/E92A mutant in Vero-AT cells (Fig. S2A).

**Figure 6.**
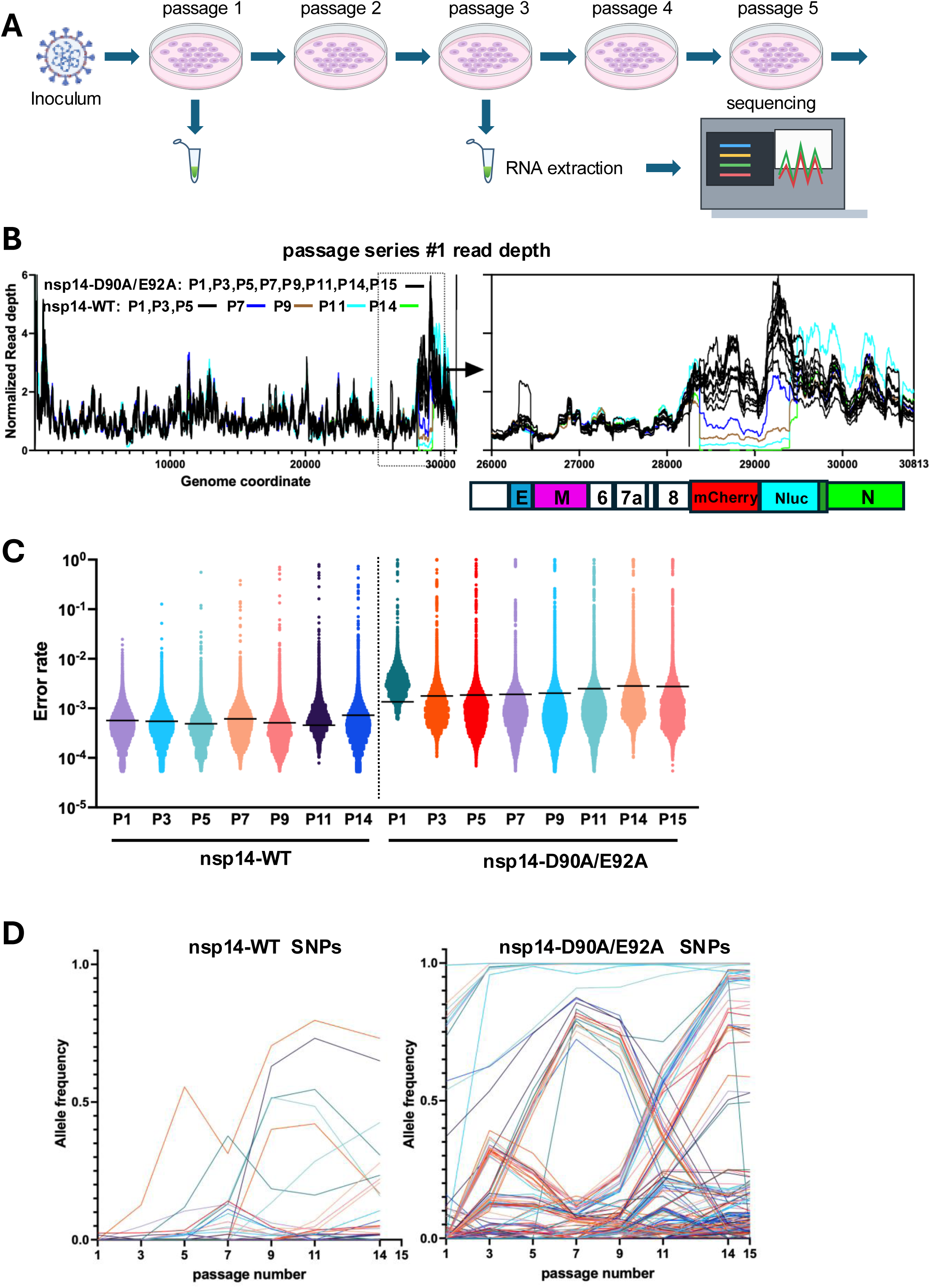
The nsp14-D90A/E92A mutation reduces replication fidelity in SARS-CoV-2. **(A)** Experimental schematic for serial passage and sequencing analysis of recombinant SARS-CoV-2. Virus was passaged in either Vero-NP or Vero-AT cells. Viral RNA was collected at passages and subjected to sequencing. **(B)** Normalized sequencing read depth across the viral genome for nsp14-WT and nsp14-D90A/E92A viruses during serial passage on Vero-NP cells. The dashed box marks the 3′ genomic region, which is shown at higher magnification in the right panel. ORFs encoded in this region are shown below the genome coordinates. **(C)** Violin plots showing the distribution of per-position error rates across the viral genome, with the mean indicated by the black line. **(D)** Allele-frequency trajectories of SNPs detected during serial passage of nsp14-WT and nsp14-D90A/E92A viruses. Each line represents a single SNP tracked across passages.

For the nsp14-D90A/E92A mutant, the engineered ExoN substitutions were stably maintained through all 15 passages. Quantification of sequence variation relative to the reference genome across passages revealed a marked increase in the apparent error frequency in the nsp14-D90A/E92A population, with mean values ranging from 0.14% to 0.28%, compared with 0.05% to 0.07% for the nsp14-WT virus (Fig. 6C, S2B and supplemental data 2). Because the baseline sequencing error rate of the Illumina platform has been reported to be approximately 0.1% [31], the values observed for the nsp14-WT virus are consistent with background sequencing error, whereas the elevated values observed for the nsp14-D90A/E92A mutant are consistent with increased viral sequence variation. Using an alternative allele-frequency cutoff of 1.5%, the nsp14-WT virus exhibited only two SNPs at passage 1 and accumulated fewer than 40 SNPs over subsequent passages (Fig. 6D, S2C and supplemental data 3). In contrast, the nsp14-D90A/E92A mutant exhibited approximately 50 SNPs at passage 1 and accumulated more than 400 SNPs by passage 14. Some SNPs reached near fixation early during passage, while others emerged transiently and were subsequently lost (Fig. 6D and S2C). Many SNPs resulted in nonsynonymous mutations in various SARS-CoV-2 ORFs (Supplemental Data 3).

Together, these results demonstrate that impairment of nsp14 ExoN activity reduces replication fidelity and substantially increases genetic diversity during SARS-CoV-2 propagation.

## Discussion

CoVs and related nidoviruses are unique among RNA viruses in encoding a proofreading ExoN that enhances replication fidelity and is thought to facilitate maintenance of unusually large RNA genomes. In this study, we show that SARS-CoV-2 can tolerate substantial impairment of nsp14 ExoN activity, but that virus recovery and replication depend on the level of ExoN function. Using a continuous 2AP-based biochemical assay, we found that mutations at the RNA-binding interface and catalytic center impair ExoN activity to different degrees. Alanine substitution of Lys9, which is positioned near the RNA substrate, had relatively modest effects on ExoN activity in vitro, likely because loss of a single substrate-contacting sidechain is not sufficient to disrupt overall RNA engagement. In contrast, charge reversal at this position, K9E, caused a severe loss of ExoN activity, likely by introducing electrostatic repulsion between the negatively charged glutamate residue and the RNA backbone, thereby interfering with productive substrate binding. Notably, this defect was more severe than that caused by alanine substitutions of the catalytic motif I residues D90 and E92. These biochemical differences were reflected in virus rescue experiments: K9A was readily recovered, D90A/E92A was recovered with delayed kinetics and reduced replication efficiency, and K9E was not recoverable. These findings support a model in which SARS-CoV-2 requires a minimal threshold of nsp14 ExoN activity for productive replication.

This threshold model helps reconcile our results with prior studies of coronavirus ExoN mutants. Motif I catalytic substitutions are tolerated in MHV and SARS-CoV, where they produce viable hypermutator viruses, but equivalent substitutions were previously reported to be lethal in MERS-CoV and SARS-CoV-2 [32]. Our recovery of SARS-CoV-2 D90A/E92A indicates that ExoN impairment is not absolutely lethal in SARS-CoV-2 and that viability likely depends on the degree of residual ExoN function, the viral genetic background, and the efficiency of virus recovery. Previous studies relied on in vitro-transcribed RNA to recover recombinant viruses, a strategy that may introduce unintended sequence variation and reduce rescue efficiency. In contrast, we employed a self-launching reverse genetics system based on a fully sequenced cDNA clone, together with cells expressing the SARS-CoV-2 N protein to enhance virus recovery. The delayed recovery and reduced growth of the D90A/E92A mutant indicate that ExoN activity contributes substantially to optimal SARS-CoV-2 replication.

Deep sequencing analysis demonstrated that the D90A/E92A mutant accumulates mutations across the genome at a substantially higher level than WT virus. The engineered mutations remained stable across passages, indicating that the mutator phenotype does not rapidly revert even during extended replication in culture. This observation parallels previous studies of ExoN mutants in other CoVs and reinforces the central role of ExoN proofreading activity in maintaining genome fidelity. The increased number of single-nucleotide polymorphisms observed during serial passage further suggests that ExoN impairment accelerates diversification of SARS-CoV-2 populations.

The ability to recover proofreading-impaired SARS-CoV-2 provides a useful system for studying how replication fidelity influences genome stability, antiviral resistance, and viral attenuation. Consistently, SARS-CoV mutants with impaired ExoN activity have previously been shown to function as live attenuated vaccines that protect aged and immunocompromised animals from lethal disease [33]. While this manuscript was in preparation, an independent study reported recovery of a SARS-CoV-2 nsp14 D90A/E92A mutant using saturation mutagenesis [34]. That study showed that this mutant has broad defects in replication, RNA synthesis, recombination, replication fidelity, nucleoside analog sensitivity, and pathogenesis. Our study is complementary but distinct in emphasis. By directly comparing defined RNA-binding and catalytic-site substitutions in parallel biochemical and reverse-genetics assays, we identify a graded relationship between residual ExoN activity and SARS-CoV-2 viability. In particular, the contrasting phenotypes of K9A, D90A/E92A, and K9E support the conclusion that SARS-CoV-2 viability depends on maintaining ExoN activity above a minimal functional threshold.

## Methods

### Cells

Vero E6 cells expressing the SARS-CoV-2 N protein (Vero-NP) were generated by lentiviral transduction of Vero E6 cells. Briefly, the pLVX-EF1α-SARS-CoV-2-N-2xStrep-IRES-Puro plasmid (Addgene #141391), together with pSPAX2 and pMD2.G (Addgene plasmids #12259 and #12260), were transfected into HEK293T cells using Lipofectamine 3000 transfection reagent. Lentiviral supernatant was collected 48 hours after transfection, centrifuged, and filtered through a 0.45 µm filter. Vero E6 cells were seeded in 6-well plates and transduced with lentivirus in the presence of 10 µg/mL polybrene, followed by centrifugation at 1500 rpm for 2 hours. After 48 hours, the cells were selected with puromycin (InvivoGen) at 5 µg/mL for 7 days. Expression of N in the cells was confirmed by Western blot using an anti-SARS-CoV N cross-reactive monoclonal antibody 1C7C7 [35].

Vero E6, HEK293T, and BHK21-ACE2 cells (gift from Guangxiang Luo, Wake Forest University) were cultured in Dulbecco’s modified Eagle’s medium (DMEM; Invitrogen) supplemented with 10% fetal bovine serum (FBS). Vero-ACE2-TMPRSS2 (Vero-AT) cells (BEI Resources) and Vero-NP cells were cultured in DMEM containing 5% FBS and 5 µg/mL puromycin.

### Bacmid mutagenesis

Recombinant SARS-CoV-2 mutants were generated using the pBAC-SARS-CoV-2/mCherryNluc or pBAC-SARS-CoV-2/Δ3a7b-Nluc bacterial artificial chromosome (BAC) clones. These BAC clones were derived from the SARS-CoV-2 USA-WA1/2020 strain (GenBank MN985325) and have been reported previously [29, 30]. Site-directed mutagenesis was performed using the *en passant* mutagenesis protocol [36]. Briefly, the bacmid was electroporated into *E. coli* strain GS1783 (gift from Greg A. Smith, Northwestern University). To introduce point mutations, primers were designed containing 30-50 bp of homology to the target region of the viral genome and ∼22 bp of homology to the kanamycin resistance cassette. The oligonucleotides used are listed in Supplementary Table 1. Oligos were synthesized and PAGE-purified (Integrated DNA Technologies). Using these primers and the pEPKan-S plasmid (gift from Greg A. Smith, Northwestern University) as template, PCR was performed to generate a linear amplicon containing the kanamycin resistance gene flanked by ∼40 bp homology arms corresponding to the target insertion site. The PCR amplicon was electroporated into *E. coli* GS1783 cells harboring the SARS-CoV-2 BAC. Recombinant clones were selected on kanamycin plates following the first Red recombination event. The kanamycin resistance cassette was subsequently removed by inducing I-SceI endonuclease cleavage with 2% L-arabinose, followed by a second Red recombination step, yielding kanamycin-sensitive clones carrying the desired point mutation. Mutant BAC DNA was purified and the presence of the engineered mutations was confirmed by whole-plasmid sequencing (Plasmidsaurus).

NSP14 mutations in the pSMART-T7-scv2-replicon [37] (BEI resources) were generated using a similar *en passant* mutagenesis strategy with the following modification. A synthesized plasmid (pAmp; Integrated DNA Technologies) containing an I-SceI recognition site and an ampicillin resistance gene was used as the PCR template instead of pEPKan-S to generate the recombination cassette, and the first recombination step was selected on ampicillin plates.

### Generating recombinant SARS-CoV-2

SARS-CoV-2 BAC DNA was purified using a QIAGEN Large-Construct kit (Qiagen) according to the manufacturer’s protocol. Five micrograms of BAC DNA was transfected into Vero-NP cells in 6-well plates using Lipofectamine 3000 transfection reagent (Invitrogen). After 4 hours, the transfection medium was replaced with DMEM containing 5% FBS. Cells were incubated at 37 °C with 5% CO₂ for 3–21 days, after which the tissue culture supernatant was collected and designated passage 0 (P0).

Subsequent passages were generated by infecting fresh Vero-NP or Vero-AT cells with 500 µL of virus from the previous passage.

### In vitro transcription, electroporation, and luciferase activity measurement

These experiments were performed essentially as previously described [37]. Briefly, pSMART-T7-scv2-replicon DNA was linearized with SwaI (New England Biolabs) and purified by phenol-chloroform extraction followed by isopropanol precipitation. The linearized DNA was used as a template for in vitro transcription using the mMessage mMachine kit (Invitrogen) according to the manufacturer’s instructions. The resulting replicon RNA was purified by phenol-chloroform extraction and isopropanol precipitation and electroporated into HEK293T cells. Remdesivir (MedChemExpress) was added to some samples at a final concentration of 10 µM. Luciferase activity at different time points after electroporation was measured using the Steady-Glo Luciferase Assay System (Promega) and quantified with a Synergy 2 plate reader (BioTek).

### Immunofluorescence assay of SARS-CoV-2–infected cells

Cells infected with recombinant SARS-CoV-2 were fixed with 4% paraformaldehyde, permeabilized with 0.1% Triton X-100, blocked with DMEM containing 10% fetal bovine serum, and stained with a rabbit monoclonal antibody against SARS-CoV-2 NP (GeneTex, GTX635679), followed by an Alexa Fluor 488-conjugated goat anti-rabbit secondary antibody (Thermo Fisher Scientific). Hoechst 33342 was added in the final step to counterstain nuclei. Fluorescence images of the entire well were acquired using a 4× objective on a Cytation 5 imaging system (BioTek).

### Plaque assay

Vero-ACE2-TMPRSS2 (Vero-AT) or BHK21-ACE2 cells were seeded in 6-well plates one day prior to infection. SARS-CoV-2 stocks were serially diluted in DMEM containing 10% FBS, and 1 mL of each dilution was added to the wells. Plates were incubated at 37 °C for 1 hour to allow viral adsorption, with gentle rocking every 15 minutes. Following adsorption, the inoculum was removed and replaced with DMEM containing 1% methylcellulose as an overlay. Plates were then incubated at 37 °C for 4 days. The overlay medium was subsequently removed, and cells were stained with 0.1% crystal violet in 20% ethanol to visualize and quantify viral plaques.

### Viral growth analysis

Confluent Vero-NP cells were infected with recombinant SARS-CoV-2 at a multiplicity of infection (MOI) of 0.005 in triplicate. After 2 hours of viral adsorption at 37 °C, the cell monolayer was washed and incubated at 37 °C with fresh medium. A total of 60 µL of culture supernatant was collected every 24 hours from day 1 to day 7 and stored at −80 °C. Nanoluciferase activity in samples from each time point was measured using the Nano-Glo Luciferase Assay System (Promega) according to the manufacturer’s instructions.

### RNA extraction and deep sequencing

Culture supernatants from serial passages of recombinant SARS-CoV-2 in Vero-NP or Vero-AT cells were processed for RNA extraction using TRIzol LS reagent (Invitrogen) according to the manufacturer’s instructions. The extracted RNA was converted into sequencing libraries and subjected to paired-end deep sequencing (100 or 150 bp reads) with NovaSeq 6000 system at the UT Health San Antonio Genomic Sequencing Core.

Sequencing reads were mapped to the reference genome using minimap2 [38]. Per-position read depth was calculated using samtools [39]. Variants were called from samtools mpileup by quantifying reference and alternate base counts at each position, followed by filtering based on sequencing quality, minimum coverage depth of 20, and a minimum alternate allele frequency of 1.5%.

### Expression and purification of SARS-CoV-2 nsp10 and nsp14 (WT and mutants)

The SARS-CoV-2 nsp10 and nsp14 bacterial expression plasmids were prepared previously [19]. To construct nsp14 mutants, we performed Quickchange site-directed mutagenesis using the WT nsp14 expression plasmid as template and appropriate DNA oligonucleotides. The final construct was confirmed by sequencing performed by Genewiz. Expression and purification of SARS-CoV-2 nsp10 and nsp14 (WT and mutants) was performed as described previously [19].

### SARS-CoV-2 ExoN-catalyzed exoribonuclease assays

Reactions contained 25 mM HEPES pH 7.5, 2 mM MgCl_2_, 1 mM TCEP, 10 mM KCl, and 50 mM NaCl and were performed at 30 °C. Typical concentrations for RNA substrate was 10 μM and enzyme was between 0.01 to 0.1 μM. Reactions were performed in 30 uL total volume in black non-binding 384-well plates. Fluorescence emission spectra were collected over time using a BioTek Synergy H1 plate reader equipped with a monochromator using an excitation wavelength of 310 nm and emission at 380 nm. dsRNA substrates were produced by annealing 100 μM RNA oligonucleotides in T_10_E_1_ [10 mM Tris pH 8.0 and 1 mM EDTA] and 50 mM NaCl in a Progene Thermocycler (Techne). Annealing reaction mixtures were heated to 90°C for 1 min and slowly cooled (5°C/min) to 10°C. Specific scaffolds are described in the figure legends. Enzymes were diluted immediately prior to use in 25 mM HEPES, pH 7.5, 500 mM NaCl, 2 mM TCEP, and 20% glycerol. SARS-CoV-2 nsp10 was pre-mixed with SARS-CoV-2 nsp14 on ice in 25 mM HEPES, pH 7.5, 500 mM NaCl, 2 mM TCEP, and 20% glycerol for 5 min prior to initiating the reaction with ExoN. The volume of enzyme added to any reaction was always less than or equal to one-tenth the total volume. The ExoN concentration refers to the nsp14 concentration and the ratio of nsp14 to nsp10 is also indicated in cases where nsp10 was in excess of nsp14. For example, 0.1 μM ExoN (1:10) refers to final concentrations of nsp14 of 0.1 μM and nsp10 at 1 μM.

### Molecular Dynamics Simulations

The starting coordinates for the nsp10-nsp14-dsRNA complex used in the molecular dynamics simulations were derived from the cryo-EM structure 7N0B [16]. The missing residues 455-464 of nsp14 were modeled using the “Model Loops” utility in ChimeraX [40]. The initial model, comprising residues 1-131 of nsp10, residues 2-523 of nsp14, dsRNA, five Zn^2+^ ions, and two Ca^2+^ ions, was subjected to all-atom molecular dynamics simulations using the AMBER software suite [41] with parameters from the amber14SB force field [42]. A second simulation was performed using a similar initial model, except that two Mg^2+^ ions replaced the Ca^2+^ ions in the first system.

All-atom MD simulations were performed in explicit water (TIP3P model [43]); a minimal distance of 20 Å between the edge of the solvent box and any protein atoms was imposed. A cutoff radius of 12 Å was used in the calculations of non-bonded interactions with periodic boundary conditions applied; particle mesh Ewald method [44, 45] was used to treat electrostatic interactions. To constrain hydrogens bonded to heavy atoms SHAKE algorithm [46] was employed. The simulations were performed by first relaxing the systems in two cycles of energy minimization; subsequently, the systems were slowly heated to 300 K using the parallel version PMEMD under NVT conditions (constant volume and temperature). Langevin dynamics [47] with collision frequency (γ=2) was used to regulate temperatures. The heated systems were then subjected to equilibration by running 100 ps of MD simulations under NPT conditions (constant pressure and temperature) with 1 fs integration time step. MD trajectories were collected over 200 ns at 1 ps interval and 2 fs integration time step. Analyses of the trajectories from MD simulations were done using CPPTRAJ program [48]. MD simulations were carried out on a multi-GPU workstation with 2x AMD EPYC 7702 64-core processor and 2x Nvidia RTX A5000.

## Acknowledgements

This work was supported by NIH grant U19AI171421 and R01AI161841. Sequencing data were generated in the UT Health San Antonio Genome Sequencing Facility (supported by NIH-NCI P30 CA054174 and 1S10OD021805-01).

## Data Information

The RNA-seq data has been deposited in the NCBI Gene Expression Omnibus under accession number GSE328556 (reviewer token: ahyfaaaadtsbdwf).

## Figure Legends

**Supplemental Figure 1.**
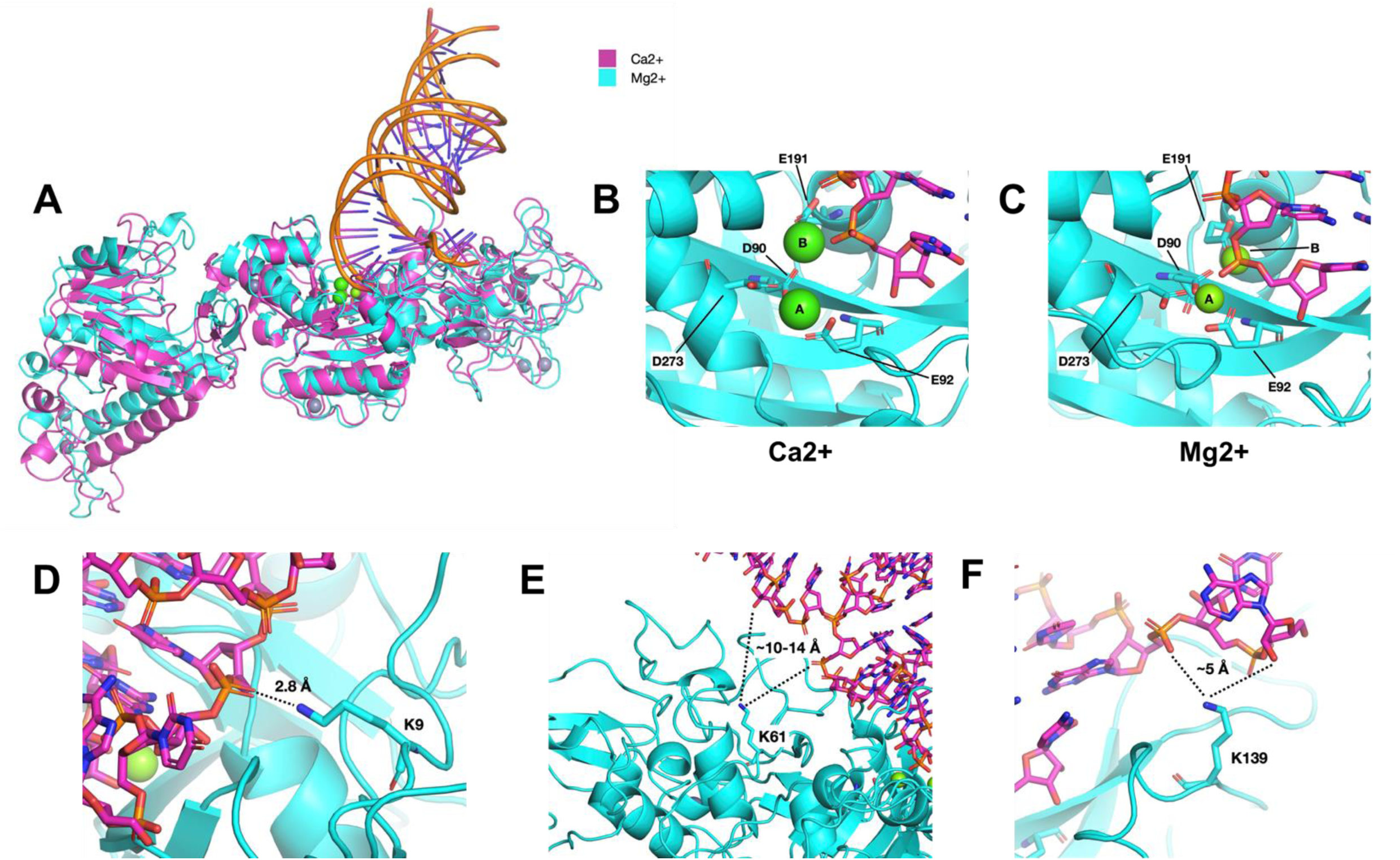
Comparison of nsp10-nsp14-dsRNA molecular dynamics (MD) simulations in the presence of Ca^2+^ or Mg^2+^. **(A)** Superimposed MD snapshots of nsp10-nsp14 and dsRNA in the presence of Ca^2+^ (magenta) or Mg^2+^ highlighting the global differences between the two simulations. Ca^2+^ and Mg^2+^ ions are shown as green spheres. Zn^2+^ ions are shown as grey spheres. The structures were superimposed using residues K9, K61, D90, E92 and K163 with an RMSD of 1.73 Å. Images were generated using the PyMOL molecular graphics program. **(B-E)** Sidechain-metal interactions between D90/E92 with Ca^2+^ **(B)** and Mg^2+^ **(C)** are highlighted. Additional coordinating residues (E191, D273) are also depicted. The relative position of the B-site coordinating ion was observed to shift resulting in a distortion of the 3’-end of the bound nucleic acid in the active site. **(D)** Residue K9 hydrogen bonds to the backbone of the templating strand of the dsRNA. The substitutions K9A or K9E (not shown) would eliminate or electrostatically repulse the interaction, respectively. **(E)** The K61 distance to dsRNA of ∼10-14 Å is shown. **(F)** K139 with backbone phosphates at the 5’-end of the template strand. Images were generated using the PyMOL molecular graphics program.

**Supplemental Figure 2.**
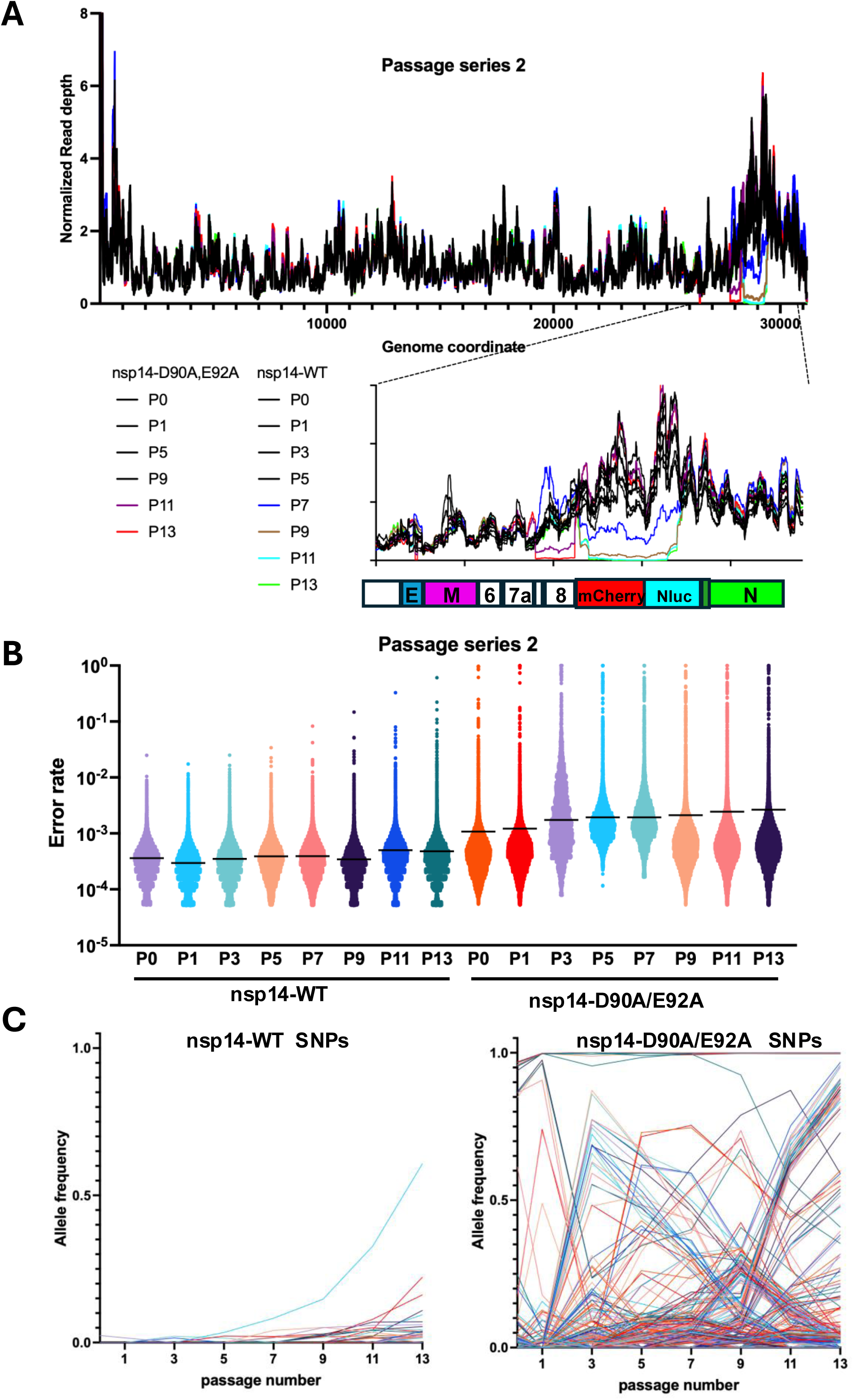
Serial passage in Vero-AT cells confirms reduced replication fidelity of the nsp14-D90A/E92A mutant. **(A)** Viruses were serially passaged as described in Figure 5, except that the passage was performed in Vero-AT cells. Viral RNA was collected at different passages and subjected to sequencing. Normalized sequencing read depth across the viral genome for nsp14-WT and nsp14-D90A/E92A viruses during serial passage is shown. The 3′ genomic region is shown at higher magnification below. ORFs encoded in this region are shown below the genome coordinates. **(B)** Violin plots showing the distribution of per-position error rates across the viral genome, with the mean indicated by the black line. **(C)** Allele-frequency trajectories of SNPs detected during serial passage of nsp14-WT and nsp14-D90A/E92A viruses. Each line represents a single SNP tracked across passages.

**Supplemental Data File 1. Sequencing read depth for all samples.** Read depth at each position of the SARS-CoV-2 genome for all sequenced samples is provided in Excel format.

**Supplemental Data File 2. Per-position variant frequencies across the SARS-CoV-2 genome.** Variant frequencies at each genomic position are shown for all viral passage samples.

**Supplemental Data File 3. SNPs and their frequencies across viral passages**. SNPs were filtered based on a minimum alternate allele frequency of 1.5%.

